# Randomized field trial of a therapeutic vaccine against *Trypanosoma cruzi* natural infection in dogs and correlates for efficacy

**DOI:** 10.1101/2025.05.12.653529

**Authors:** Jorge A. Calderón-Quintal, Christian F. Teh-Poot, Landy M. Pech Pisté, Pedro P. Martinez-Vega, Victor Dzul-Huchim, Felipe Torres-Acosta, Etienne Waleckx, Liliana Villanueva-Lizama, Jaime Ortega-Lopez, Claudia Herrera, Eric Dumonteil, Julio Vladimir Cruz-Chan

## Abstract

**Background and Aims:** Chagas disease, caused by *Trypanosoma cruzi*, is a vector-borne parasitic disease, with dogs acting as a major domestic host of the parasite. An immunotherapeutic vaccine would be an excellent tool to treat infections and prevent chronic cardiac disease in this host. Building on previous pre-clinical studies, we performed here the first randomized field trial of a vaccine against *T. cruzi* among client-owned dogs with natural infections.

**Methods:** A total of 31 dogs with *T. cruzi* infection with diverse parasite strains were enrolled and received three doses of a vaccine composed of Tc24-C4 and TSA1-C4 recombinant proteins with MPLA (N=16) or saline control (N=15) and followed for up to six months to assess efficacy.

**Results:** Blood parasite burden and electrocardiographic (ECG) recordings as primary outcomes showed that therapeutic vaccination led to a significant decrease in parasite burden, prevented/stopped cardiac alterations and was safe. This clinical benefit was mediated by major changes in T cell activation and T cell receptor (TCR) repertoire, while antibody responses were minimally affected. In addition, vaccination also reprogrammed the ongoing trained immunity to reduce inflammation, suggesting a complex interplay between innate and T cells in its mechanism of action.

**Conclusions:** These results provide a strong support for the further development of a veterinary vaccine based on these antigens as well as a human therapeutic vaccine to prevent the progression of chronic cardiac disease from *T. cruzi* infection.

**Structured graphical abstract:** *Key question:* Can an immunotherapeutic vaccine against *Trypanosoma cruzi* control an ongoing infection and prevent the progression of chronic cardiac disease in naturally infected dogs?

*Key finding:* Therapeutic vaccination of chronically infected dogs led to a reduced blood parasite burden and prevented the progression of chronic cardiac disease (Primary outcomes). Correlates of vaccine efficacy included major changes in T cell repertoire/activation and changes in innate immunity to reduce inflammation.

*Take home message:* Immunotherapeutic vaccination of dogs with natural infection with *T. cruzi* was safe and effective to control an ongoing infection with a broad diversity of parasite strains and prevented the progression of chronic cardiac disease. 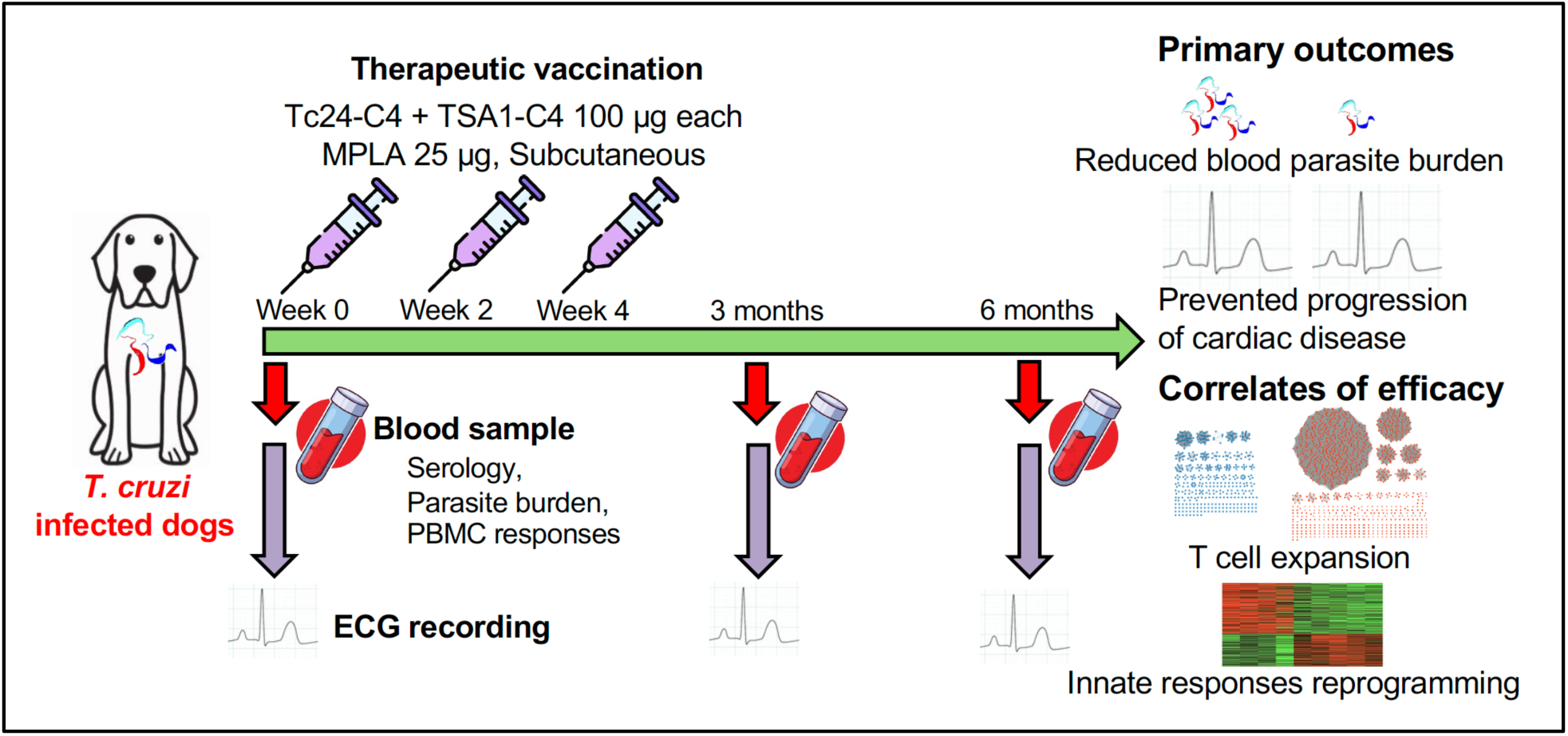

*Translational perspective:* Current drug treatments for Chagas disease have major limitations due to significant adverse effects and a limited efficacy as chronic cardiac disease develops. A veterinary vaccine for dogs, a major domestic host of the parasite, would help reduce domestic transmission and improve dog health, as well as provide support for the development of a human immunotherapeutic vaccine to prevent the progression of chronic Chagasic cardiomyopathy.

## INTRODUCTION

Chagas disease, caused by the protozoan parasite *Trypanosoma cruzi*, is a vector-borne neglected tropical disease endemic in most of the Americas. While the parasite circulates mostly as a zoonotic agent among wild vectors and mammals, the intrusion and domiciliation of some vector species can result in domestic parasite transmission. Thus, there are at least 6 million cases in Latin America and 300,000 cases in the US ^1,2^, contributing to a global annual burden of $627.46 million in health-care costs and 806,170 disability-adjusted life-years (DALYs) ^3^.

Dogs are a major domestic host of *T. cruzi* parasites, increasing the risk for human infection and they can serve as a sentinel for domestic transmission cycle ^4–7^. In addition, dogs may present clinical signs of disease with similarities to human patients with Chagas disease including cardiac alterations, leading to comparable morbidity and mortality. For example, naturally infected dogs can present ventricular arrhythmias, atrioventricular block, sinusal block, right bundle branch block, QRS complex alterations, or supraventricular premature contractions ^8–12^.

Current drug treatments with benznidazole and nifurtimox have severe limitations due to a low efficacy during the chronic phase *of T. cruzi* infection and important adverse effects that complicate treatment completion ^13–16^. Thus, an immunotherapeutic vaccine would provide an excellent alternative/complement to treat infected hosts and prevent or at least delay the development of cardiac manifestations by decreasing parasite burden and reducing inflammation and fibrosis ^17–20^. A vaccine candidate based on Tc24 and TSA1 parasite antigens has been extensively tested in different formulations in mouse models and showed promise in controlling experimental infections with *T. cruzi* ^21–25^. In terms of mechanisms of action, current understanding is that therapeutic vaccination leads to a reorientation of the ongoing immune response toward a balanced Th1/Th2/Th17 profile with the strengthening of CD8^+^ T cell responses, allowing for a better parasite control ^26–28^. Additional studies indicate that vaccination may be combined with benznidazole treatment for increased therapeutic efficacy ^29–31^.

A veterinary vaccine would be a key tool for improved control of *T. cruzi* ^32^ and these antigens formulated as DNA vaccines were effective as preventive or therapeutic vaccines in dogs with experimental *T. cruzi* infection ^33^. Although both antigens are highly conserved among *T. cruzi* strains ^34,35^, it remains unclear how this vaccine candidate may perform in the context of natural infections, which involve a broad diversity of strains ^36–38^. Therefore, we evaluated here the therapeutic efficacy of this vaccine candidate, formulated as a recombinant protein vaccine with monophosphoryl lipid A (MPLA) as adjuvant ^22,25,39^, in a first field trial among client-owned dogs with natural *T. cruzi* infection. The trial was performed in a rural endemic area in southern Mexico, where around 27% of dogs are infected with a broad diversity of *T. cruzi* strains ^37^. As primary outcome, blood parasite burden and cardiac function were evaluated following vaccine treatment. The correlates of vaccine efficacy were also investigated by evaluating the immune response and the transcriptomic profile of peripheral blood mononuclear cells (PBMCs). Our results provide a strong rationale for the further development of a veterinary vaccine against *T. cruzi* based on these antigens as well as for a human therapeutic vaccine ^17,18^.

## MATERIALS AND METHODS

### Ethics statement

All work was approved by the Institutional Bioethics Committee of the “Centro de Investigaciones Regionales Dr, Hideyo Noguchi”, Universidad Autónoma de Yucatán (Reference # CEI-05-2021) and was performed in compliance with NOM-062-ZOO-1999. Informed consent was obtained from dog owners prior to enrolment of their dogs in the study.

### Study design

A cohort of 31 client-owned dogs previously diagnosed with natural *T. cruzi* infection were included in this study. They were assigned to a treated or placebo group by stratified randomization to ensure an even distribution of sex and age between the two groups. Half of the dogs (N=16 dogs) were treated with three doses of a therapeutic vaccine against *T. cruzi* by subcutaneous injections at two-week intervals (0, 2 and 4 weeks). The other half of the dogs (N=15) received saline solution as placebo control. A minimal sample size of 13 dogs/group was determined to be able to detect a significant decrease in parasite burden of at least 45% in the vaccine treated groups with a power of 80% and our sample size exceeded this minimum. Dogs were then followed at three and six months after immunization, for general physical examination as part of safety evaluation, for the collection of blood samples for parasitological and immunological analysis, and cardiac functional assessment by ECGs. A wellness check was performed at 12 months after immunization to assess survival but no examination or sample collection was performed at that time. The primary outcomes were blood parasite burden and ECG functional profiles to evaluate therapeutic efficacy at three and six months, secondary outcomes included immune response and PBMC transcriptomic profile following therapeutic vaccination, to identify the correlates of vaccine effects. Personnel performing laboratory assays and ECG analysis were blinded to the group allocation of individual dogs. The study followed the ARRIVE Guidelines for animal studies ^40^.

### Animals

Client-owned dogs from the rural village of Sudzal, Yucatan, Mexico, with *T. cruzi* infection confirmed by a positive parasite PCR were included in the study ^37^. Dogs were infected with various mixtures of parasite DTUs, although TcI predominated ^37^. Both male and female dogs were enrolled, all mixed breed, aged 3-4 years old (Table 1). All dogs were dewormed with Vermi Plus (AlphaChem, Mex) 1-2 weeks before the first immunization. Dams developing a pregnancy were excluded from the study. Ten dogs without *T. cruzi* infection from the same village were used as controls for ECG recordings and negative blood samples. All animals remained under the care of their owners in their usual environment during the entire study.

**Table 1.** Characteristics of enrolled dogs.

|  | Saline control<br>(N=15) | Vaccine<br>(N=16) | P value |
| --- | --- | --- | --- |
| Sex (Male/Female) | 11/4 | 11/5 | 0.8 |
| Age (years) | 3.4 ± 0.8 | 3.9 ± 0.8 | 0.6 |
| Parasite burden (para. Eq./ml blood) | 10.1 ± 6.1 | 7.9 ± 6.1 | 0.8 |
| Parasite DTUs (N (%)) |  |  |  |
| Tcl only | 6 (40) | 3 (19) | 0.2 |
| Mixed DTUs* | 4 (27) | 9 (56) |  |
| Unknown | 5 (33) | 4 (25) |  |
| ECG alterations (N (%)) |  |  |  |
| None | 7 (47) | 10 (62) | 0.2 |
| Electrical left axis deviation | 5 (33) | 5 (31) |  |
| First-degree atrioventricular block | 2 (13) | 0 (0) |  |
| Bradycardia | 0 (0) | 1 (6) |  |

\* Mixed DTUs are mixtures of Tcl with TcII, TcIV, TcV and/or TcVI in variable proportions.

### Vaccine and vaccination

The recombinant proteins Tc24-C4 and TSA1-C4 were produced as described before ^22,25,41^ and stored at −80°C until used. Purity was >95% for each protein and integrity was tested by SDS-PAGE electrophoresis prior use. Vaccinated dogs received three doses of 100 µg of each recombinant TSA-1-C4 and Tc24-C4 proteins with 25 µg of MPLA adjuvant in 500 µl of PBS by subcutaneous injections at 2 weeks interval (week 0, 2 and 4). Animals in the placebo group received three doses of 500 µl of saline solution. The dogs were monitored for 30 min after administration of each dose to monitor for severe immediate, adverse events. Dog owners were contacted within 48 h to record any adverse events that occurred within that time.

### Blood parasite burden

Blood samples were collected from the cephalic vein with 0.05% EDTA as anticoagulant for the preparation of plasma and PBMCs that were cryopreserved. DNA was extracted using Wizard genomic purification kit (Promega Inc). The parasite burden was determined by a standardized quantitative real-time (qPCR) assay ^42^. Briefly, this TaqMan duplex assay targets *T. cruzi* satellite DNA (primers cruzi1/cruzi2 and probe cruzi3) and RNAse P gene as an internal amplification control. It also includes Uracil-DNA Glycosylase (UDG) as a carry-over contamination control. Cycling conditions consisted of 2 min at 50 °C and 10 min at 95 °C, followed by 45 cycles of 95 °C for 15 s and 58 °C for 1 min. Positive parasite and water negative controls were included alongside samples with each PCR run and all samples were run in duplicate. Cq values were converted to parasitic load (parasite equivalents/mL) based on a standard curve.

### Humoral and cellular immune response

Antigen-specific antibody responses were measured at different time points in plasma samples through indirect ELISA as before ^37^. We measured anti-*T. cruzi* IgG antibodies using a parasite soluble antigen extract, as well as IgG against Tc24-C4 and TSA1-C4 vaccine antigens. For cellular immune responses, peripheric blood mononuclear cells (PBMCs) were purified and stimulated *in vitro* with either 20 µg *T. cruzi* soluble antigen, 25 µg Tc24-C4 or 25 µg TSA1-C4 antigens for 24 h prior to staining for CD3, CD4 and CD8 surface molecules, and analysis by flow cytometry as described before ^37^.

### Electrocardiography

Electrocardiograms (ECGs) were recorded on unsedated dogs in right lateral recumbency, with electrode clips on the medial knee and elbows, using an electrocardiograph ECG100G-VET with a paper speed of 50 mm/s and 10mm/mV. The recordings were obtained in Lead II and the main waves and interval were measured, including P wave, PR interval, QRS complex, QT interval, ST segment, T wave and mean electrical axis ^37^.

### PBMC transcriptomic analysis

Total RNA was purified from unstimulated PBMCs collected at the three months time point, using E.Z.N.A.® Total RNA Kit (Omega Bio-Tek), and stored at room temperature in Gen-Tegra RNA tubes until used. Illumina libraries were prepared from 400 ng of RNA/sample and sequenced on a NovaSeq platform to give an average 20 million reads/sample. Raw sequences have been deposited in NCBI SRA database, BioProject #PRJNA1248531, Biosamples SAMN47860485-SAMN47860493. Sequences were mapped to the dog reference genome (NCBI accession GCF_011100685.1) with Geneious Prime 2024 RNA mapper, and raw read counts were generated. Differential expression of protein coding genes as well as that of long non coding RNAs (lncRNAs) was assessed with DESeq2 ^43^ with the iDEP 2.0 platform ^44^, using FDR-adjusted p values <0.05 and a fold change of ± 1.5 for statistical significance. Pathway analysis was performed on differentially expressed genes using Gene Ontology (GO) enriched pathways in ShinyGO ^45^. Additional pathway enrichment analysis was performed with Gene Set Enrichment Analysis (GSEA) 4.3.2.^46^. Annotated gene sets from the Hallmark molecular signature database ^47^ and the blood transcriptome module (BTM) ^48^ were used for these analyses. The statistical significance of pathways differentially regulated by vaccine treatment was based on FDR-adjusted P values.

For immune repertoire analysis, RNA sequence reads were mapped to the CDR3 variable region of IgG heavy chain and of the T cell receptor beta chain, respectively ^49^. Mapped sequences were then analyzed through the IMGT/HighV-QUEST version 1.9.5 platform ^50^ and IMGT/StatClonotype R package ^51^ to determine V, D, J gene usage for both TCR and IGH CDR3 domains. CDR3 sequence length and composition were compared to assess the overlap of dog CDR3 sequences among individual dogs and treatment groups. CDR3 sequence clonotypes were considered public when found in more than one individual dog. Richness, Simpson and Shannon diversity indices ^52^ were calculated to assess global changes in repertoire diversity. Networks of CDR3 repertoires were elaborated in Cytoscape, based on sequence similarity analysis in EFI as before ^49^.

### Dog leukocyte antigen (DLA) typing

Allele types for DLA class I (DLA-79, DLA-88) and class II (DLA-DRB1, DLA-DQB1 and DLADQA1) were identified by mapping RNA sequence reads to the respective reference genes, and reads were de-novo assembled to produce allele sequences. These were then compared with a curated database of dog DLA alleles from the IDP-MHC repository using BLAST and assembled alleles were considered a match when showing a >99% sequence identify with a known DLA-allele ^49^.

### Statistical analysis

The results were presented as individual point graphs, or as mean ± SEM. The normality of the data was evaluated using Kolmogorov-Smirnov test. Continuous variables were compared between the vaccine-treated and control group by Student’s t test or Wilcoxon as corresponded, or by ANOVA followed by Tukey post-hoc test. X^2^ tests were used for the comparison of categorical variables. The ECG parameters were also integrated into a multivariate analysis, and a Linear Discriminant Analysis (LDA) was used to evaluate the effect of treatment on ECG patterns ^37,53^. The statistical significance of differences was assessed by permutational Multivariate Analysis of Variance (PERMANOVA) with Bonferroni correction to adjust P values for multiple comparations.

## RESULTS

### Dog characteristics

A total of 31 dogs with natural *T. cruzi* infection were enrolled in the study. They were aged 3-4 years old, mostly males, and an average blood parasite burden of 8-10 equivalent parasites/ml (Table 1). Infection consisted of mostly TcI parasite strains, although many dogs had infections that included mixtures of TcII, TcIV, TcV and TcVI parasite discrete typing units (DTUs) in variable proportions. Most dogs had normal electrocardiographic (ECG) recordings, some had minor alterations such as a left axis deviation, two dogs presented a first degree atrioventricular block, and one bradycardia (Table 1). Overall, these characteristics were distributed evenly between the two groups of dogs, and none of the differences reached statistical significance.

Dog DLA was also typed in four control and five vaccinated dogs for two class I (DLA-79 and DLA-88), and three class II (DRB1, DQA1 and DQB1) alleles, showing no significant differences in allele frequencies between groups, except for DLA-88, likely due to the small number of dogs typed (Supplementary Table 1).

### Vaccine safety

A total of 16 dogs were then treated with three doses of therapeutic vaccine based on Tc24-C4 and TSA1-C4 recombinant proteins formulated with MPLA via subcutaneous injections, at 2 weeks interval, and 15 dogs received saline as placebo control. No acute adverse reactions were detected during the 30 min of observation following each vaccine dose. No longer term adverse effects were reported by dog owners, and all dogs presented normal behavior and activities. During follow-up at three months, there were 11 dogs (73%) remaining in the control group as two were withdrawn from the study due to pregnancy, one suffered from an accidental death, and one died likely from distemper (Supplementary Table 1). No loss to follow-up occurred in the vaccine group at this time (N=16, 100%). At the six months follow-up, an additional four control dogs were lost, one from accidental death, one owner discontinued the participation of their dog in the study, one died from distemper, and one had no information regarding loss. In the vaccine group, one dog was withdrawn from the study due to pregnancy. Thus, success of follow-up at six months was 47% for control dogs, and 94% for vaccinated dogs (Supplementary Table 1). We further reached out to dog owners at 12 months to inquire about their dogs. Five of the control dogs were alive (33%), and two were lost to follow up, while for vaccine treated dogs 11 were confirmed to be alive (69%). One vaccine-treated dog had died from accidental death, two suffered potential distemper, and one unknown death occurred (Supplementary Table 2). Overall, these data suggest that the vaccine was safe and well tolerated.

### Therapeutic vaccine efficacy

Therapeutic vaccine efficacy was evaluated by measuring blood parasite burden and cardiac function by electrocardiographic (ECG) recordings as primary outcomes at three and six months after the first vaccine dose. As expected, control dogs had a steady blood parasite burden during the six months of follow-up. On the other hand, vaccine treated dogs presented a significant decrease in blood parasite burden at six months (Figure 1A, t=2.28, P=0.035). Furthermore, 5/13 vaccine treated dogs were PCR negative for *T. cruzi* at that time, while all control dogs remained PCR positive (X^2^=5.17, d.f.=1, P=0.023). Among vaccine treated dogs, those that became PCR negative were infected with mixtures of *T. cruzi* DTUs (5/5, 100%), while those that remained PCR positive were predominantly infected with TcI only (3/8, 37.5%) and only 2/8 (25%) were infected with a mixture of DTUs (and 3/8, 37.5% with unknown parasite genotype)(X^2^=8.9, d.f.=2, P=0.011), suggesting that vaccine efficacy may be higher against mixtures of parasite DTUs.

**Figure 1.**
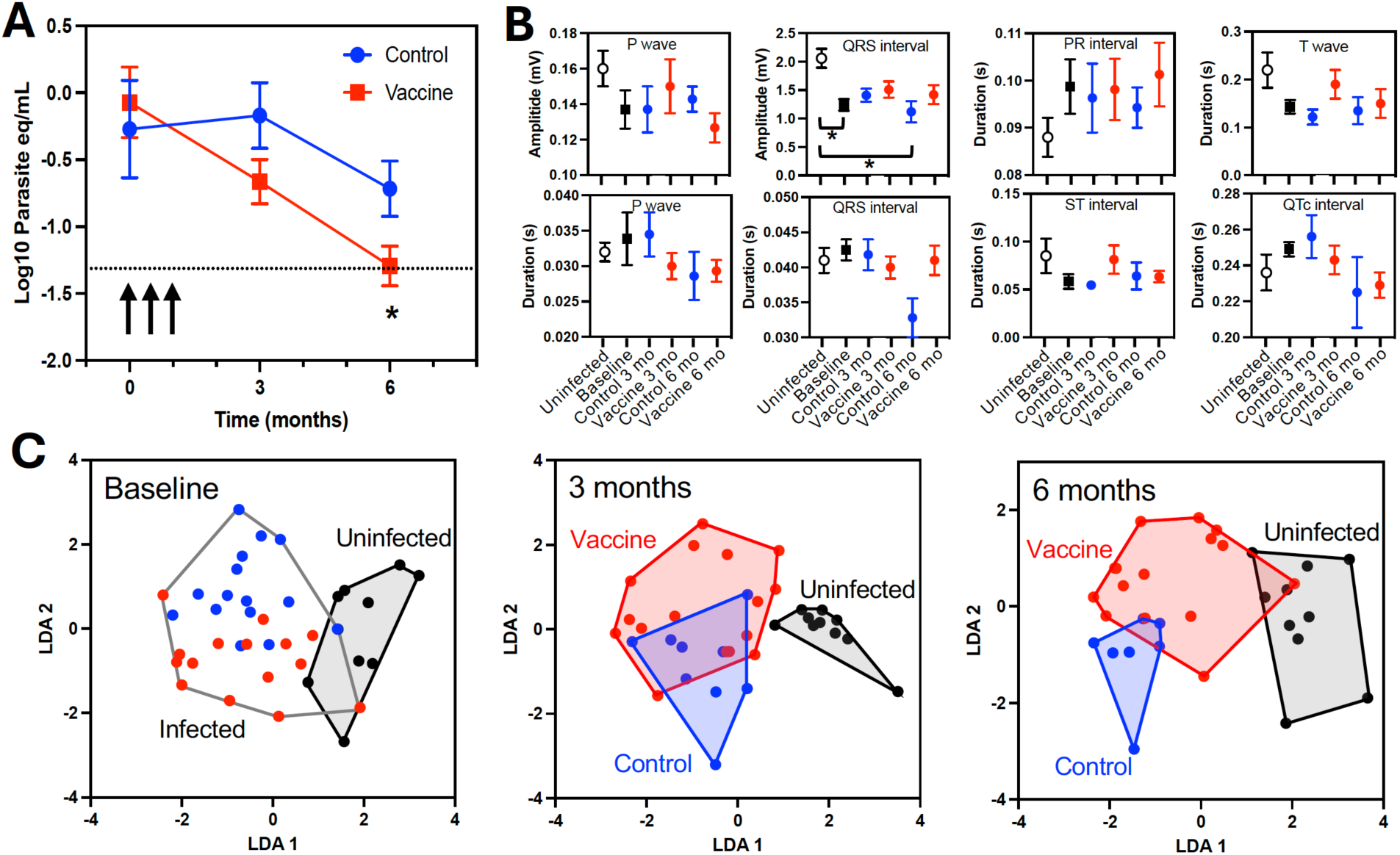
Parasitological and cardiac function primary outcomes of vaccine treatment efficacy. (**A**) Blood *T. cruzi* parasite burden measured by qPCR following vaccine treatment (arrows). There was a significant decrease in parasite burden at six months post-vaccination (*, t=2.28, P=0.035). The dotted horizontal line indicates the threshold for PCR negative samples. (**B**) Individual ECG parameters of dogs during follow-up, including P wave amplitude, P wave duration, QRS amplitude, QRS duration, PR, ST and QTc intervals, and T wave duration, measured at baseline, three- and six-months post-vaccination. Data are presented as mean ± SEM of 7-16 dogs/group and * indicates a significant difference between groups by Tukey post-hoc test (P<0.05). (**C**) LDA of ECG parameters at the indicated time-points post-vaccination. Each dot represents an individual dog, which are color-coded for uninfected (Black), vaccine-treated (Red) and untreated controls (Blue). At baseline, there was a significant difference in the ECG profile of *T. cruzi* infected dogs compared to uninfected dogs (PERMANOVA F=1.96, P=0.001). At the 3 months time-point, the ECG profile of vaccine-treated and control dogs remained significantly different from uninfected dogs (F=1.54, P=0.024 and P=0.002, respectively), and there was no significant difference between these two experimental groups (P=0.76). At 6 months, only the ECG profile of untreated control dogs was significantly different from that of uninfected dogs (F=1.66, P=0.005), while that of vaccine treated dogs did not reach statistical significance (P=0.074).

Assessment of cardiac function through ECG recordings indicated that no major arrythmias developed in any of the dogs during the six months of follow-up. In addition, no significant differences in ECG waves and intervals were detected among vaccinated dogs, control dogs, and uninfected dogs, except for QRS voltage amplitude, which was significantly decreased in infected dogs at baseline compared to uninfected dogs, and this difference with uninfected dogs was maintained at six months follow-up for control dogs, but not for vaccinated dogs (Figure 1B), suggesting potential improvement in cardiac function in response to vaccine treatment. Multivariate analysis of the ECG parameters revealed further differences in ECG profiles among groups. Indeed, at baseline, infected dogs presented altered ECG profiles compared to uninfected dogs (PERMANOVA F=1.96, P=0.001), compatible with early onset cardiac alterations, and this difference was maintained at three months post-treatment in both the vaccinated and control group (F=1.54, P=0.024 and P=0.002, respectively, Figure 1C). At six months post-treatment, the ECG profile of untreated control dogs was significantly different from that of uninfected dogs and appeared shifted further away (F=1.66, P=0.005), suggesting cardiac disease progression, while the ECG profile of vaccine-treated dogs remained close to that of uninfected controls and the differences did not reach statistical significance (P=0.074). Together, these data indicated a clear decrease in blood parasite burden following therapeutic vaccination and show that the progression of cardiac alterations was stopped/delayed for at least six months.

### Immune response and correlates of vaccine efficacy

To assess the mechanisms underlying vaccine efficacy, humoral and cellular responses were measured. Remarkably, there were no significant differences in IgG levels against *T. cruzi* soluble antigen, or vaccine antigens Tc24-C4 and TSA1-C4 over time in either control or vaccine-treated dogs (Figure 2A). On the other hand, a significant increase in TSA1-C4-specific CD8^+^ T cells was detected at three months following the first vaccine dose (t=3.01, P=0.017), although CD8^+^ cell response to Tc24-C4 or to *T. cruzi* soluble antigen were not detected (Figure 2B). Thus, vaccine treatment seemed to promote CD8^+^ T cell responses rather than antibody responses.

**Figure 2.**
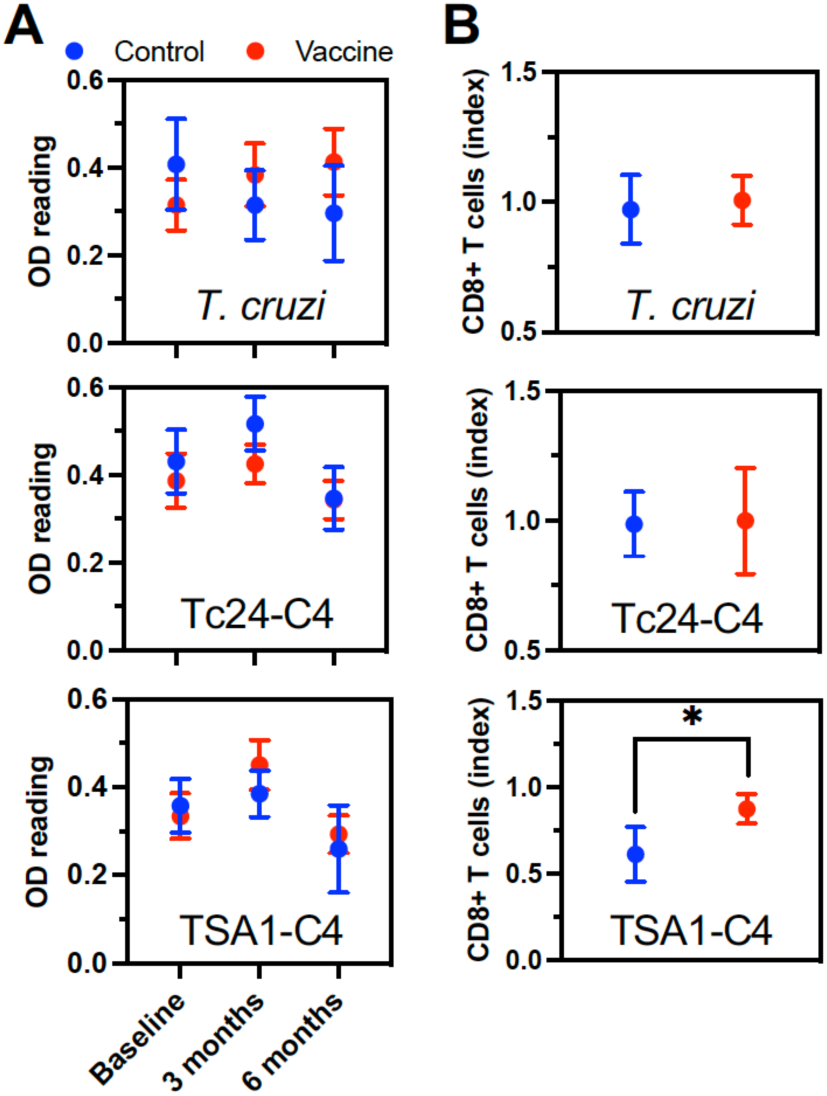
Humoral and cellular immune response of vaccine-treated dogs. (**A**) IgG against *T. cruzi* soluble antigen (top), Tc24-C4 (middle) and TSA1-C4 antigens (bottom), measured by ELISA at the indicated time points. Data are presented as the mean ± SEM of 7-16 dogs per group. There were no significant differences in antibody levels between groups or over time. (**B**) Antigen-specific CD8+ T cell recall response, measured by flow cytometry at three months after the first vaccine dose, for *T. cruzi* soluble antigen (top), Tc24-C4 (middle) and TSA1-C4 antigens (bottom). There was a significant recall of TSA1-C4-specific CD8+ T cells (*, t=3.01, P=0.017).

To further understand therapeutic vaccine effects, a transcriptomic analysis of PBMCs from a subset of dogs was performed at three months after the first vaccine dose. A total of 1263 genes were found to be differentially expressed between PBMCs from untreated controls and vaccinated dogs, corresponding to 889 downregulated and 374 upregulated genes (Figure 3A and B). PCA of the differentially expressed genes showed a marked difference in expression profile between cells from control and vaccinated dogs (Figure 3C), evidencing long-term effects of the therapeutic vaccination on PBMCs. Pathway enrichment analysis indicated that upregulated genes participated in transcriptomic remodeling and in multiple metabolic pathways including nucleic acid, aromatic compound, or nitrogen compound metabolism (Figure 3D). On the other hand, downregulated genes were involved in multiple immune-related processes suggesting broad changes in the immune response. In particular, the inflammatory and innate response pathways were downregulated (Figure 3E). Deconvolution of PBMC cell type composition based on transcriptomic profiles further indicated important changes. Indeed, the proportion of memory B cells was significantly decreased in PBMCs from vaccinated dogs, while that of CD4^+^ and CD8^+^ T cells was increased (Figure 3F), again suggesting that therapeutic vaccination promoted T cell activation rather than B cell responses. In addition to its effects on the adaptive immune response, vaccination led to a reduction in NK cells and monocytes populations (Figure 3F), suggesting a long-term modulation of the innate immune response.

**Figure 3.**
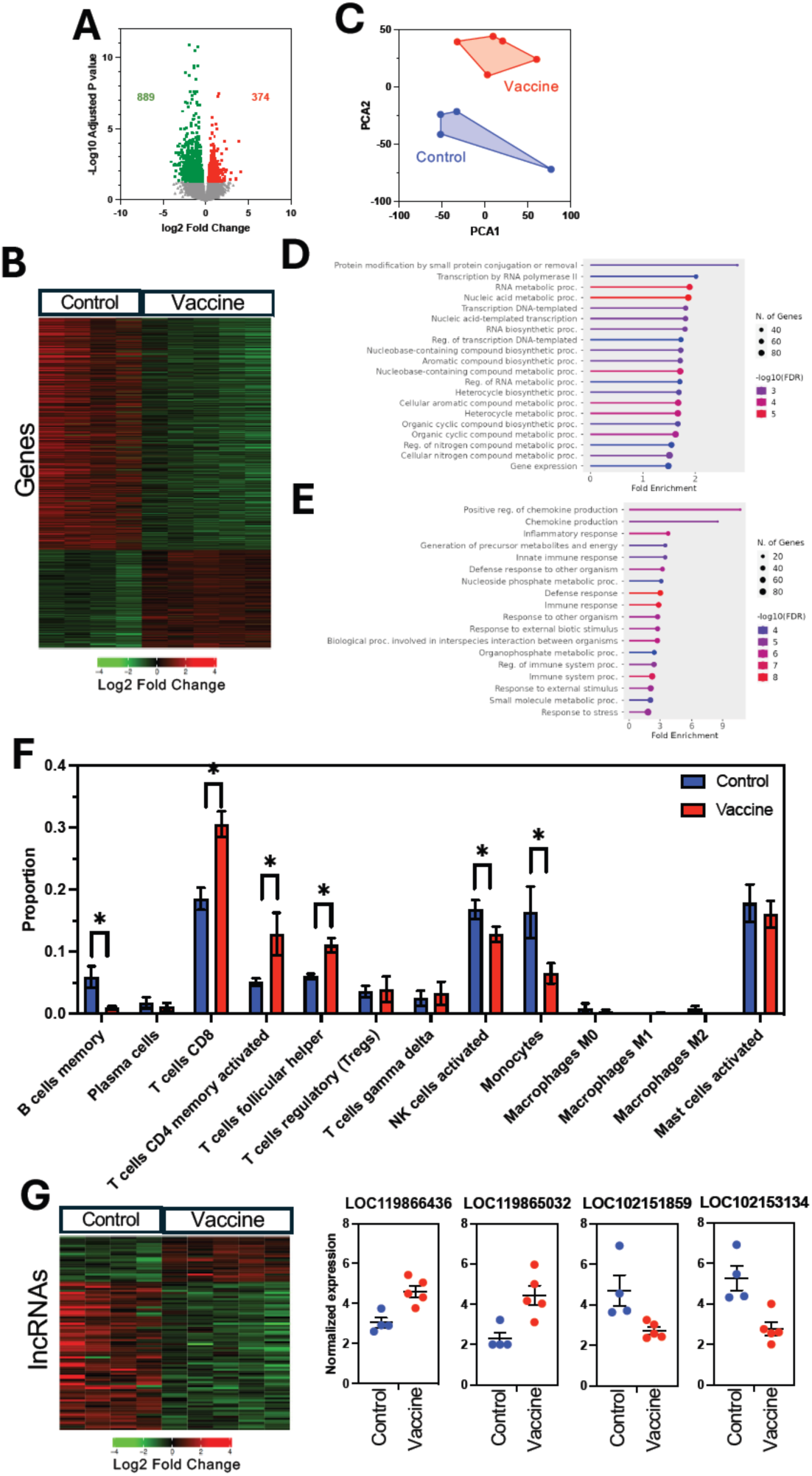
Differentially expressed genes in PBMCs from vaccine-treated dogs. (**A**) Volcano plot of differentially expressed genes indicating 889 significantly downregulated and 374 up-regulated genes in vaccinated dogs compared to control dogs (FDR adjusted P<0.05). (**B**) Heatmap of differentially expressed genes of PBMCs from vaccinated and control dogs. (**C**) Principal component analysis of differentially expressed genes. Up-regulated (**D**) and downregulated (**E**) pathways associated with the differentially expressed genes. (**F**) Deconvolution of PBMC cell type composition based on transcriptomic data. * Indicates a significant difference between cell proportion of vaccinated and control dogs presented as mean ± SEM. (**G**) Heatmap and dot plots of differentially expressed lncRNAs in PBMCs from control and vaccine treated (FDR adjusted P<0.05). All data are from four control and five vaccinated dogs.

Interestingly, further analysis allowed to detect 94 differentially expressed lncRNAs following vaccination, as 22 lncRNAs were upregulated and 72 were downregulated (Figure 3G). While no specific function could be associated with these lncRNAs, they may be involved in the modulation of some of the genes and pathways affected by therapeutic vaccination. They may also serve as potential biomarkers of vaccine therapeutic effects.

Changes in the immune response following therapeutic vaccination were further supported by Gene Set Enrichment Analysis (GSEA) (Figure 4), which revealed that multiple pathways associated were significantly altered by therapeutic vaccination. As expected, adaptive immunity pathways associated with B and T cells and IFNγ response were modulated by vaccination (Figure 4). In addition, multiple pathways associated with the innate immune response were also modulated by vaccination. These included pathways associated with monocytes, neutrophils, and dendritic cells, as well as TLR inflammatory signaling and the inflammatory response pathways (Figure 4). Additional pathways for complement, reactive oxygen species, antimicrobial peptides, and interactions between lymphoid and non-lymphoid cells were also affected by vaccination, together with glycolysis and to a lesser extent fatty acid metabolism (Supplementary Figure 1), consistent with metabolic changes in PBMCs.

**Figure 4.**
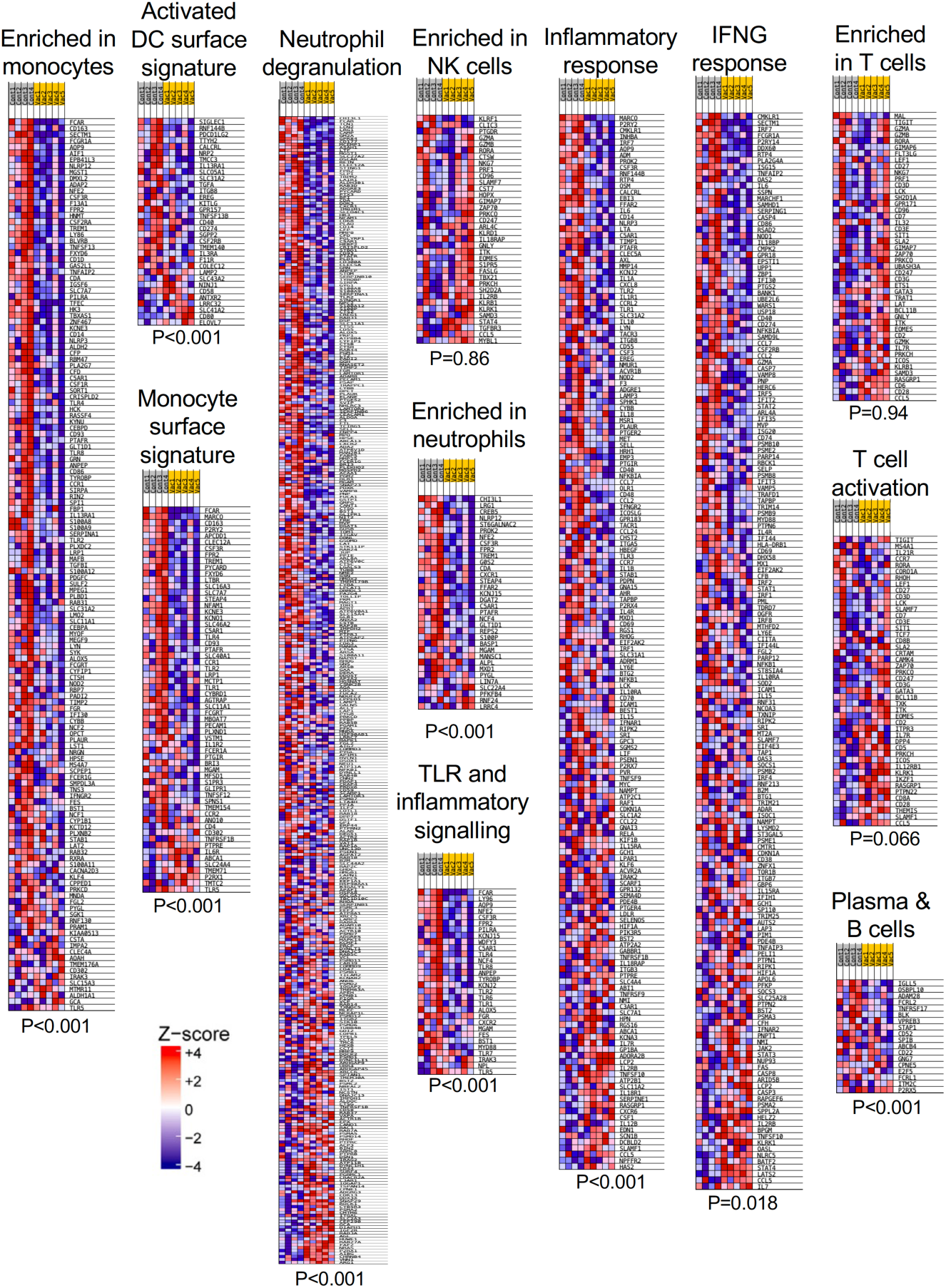
GSEA pathways in vaccine-treated and control dogs. Heatmaps of differential gene expression profiles from selected GSEA pathways among vaccine-treated (Vac) and control dogs (Cont) with *T. cruzi* infection. The statistical significance of the pathways is indicated at the bottom of each heatmap (FDR-adjusted P value).

We then focused in more details on IgG and TCR repertoires, to assess the breadth of the adaptive response among *T. cruzi* infected dogs with and without therapeutic vaccination based on expressed CDR3 sequence diversity within IgGH and TCRB molecules. Analysis of IgGH CDR3 repertoire indicated a rather limited effect of therapeutic vaccination (Figure 5A and B), although a significant decrease in richness was detected (Figure 5B), but no significant changes in Shannon or Simpson diversity indices: Thus, there was a somewhat less diverse antibody repertoire in vaccine-treated dogs, in agreement with the downregulation of B cells detected above. Importantly, there was no overlap in IgGH CDR3 repertoire among dogs and between groups, with only 1/2185 (0.05%) public IgGH CDR3 among control dogs, and 1/1368 (0.07%) among vaccine-treated dogs (Figure 5C), suggesting that the antibody response of individual dogs targeted very different parasite epitopes/antigens.

**Figure 5.**
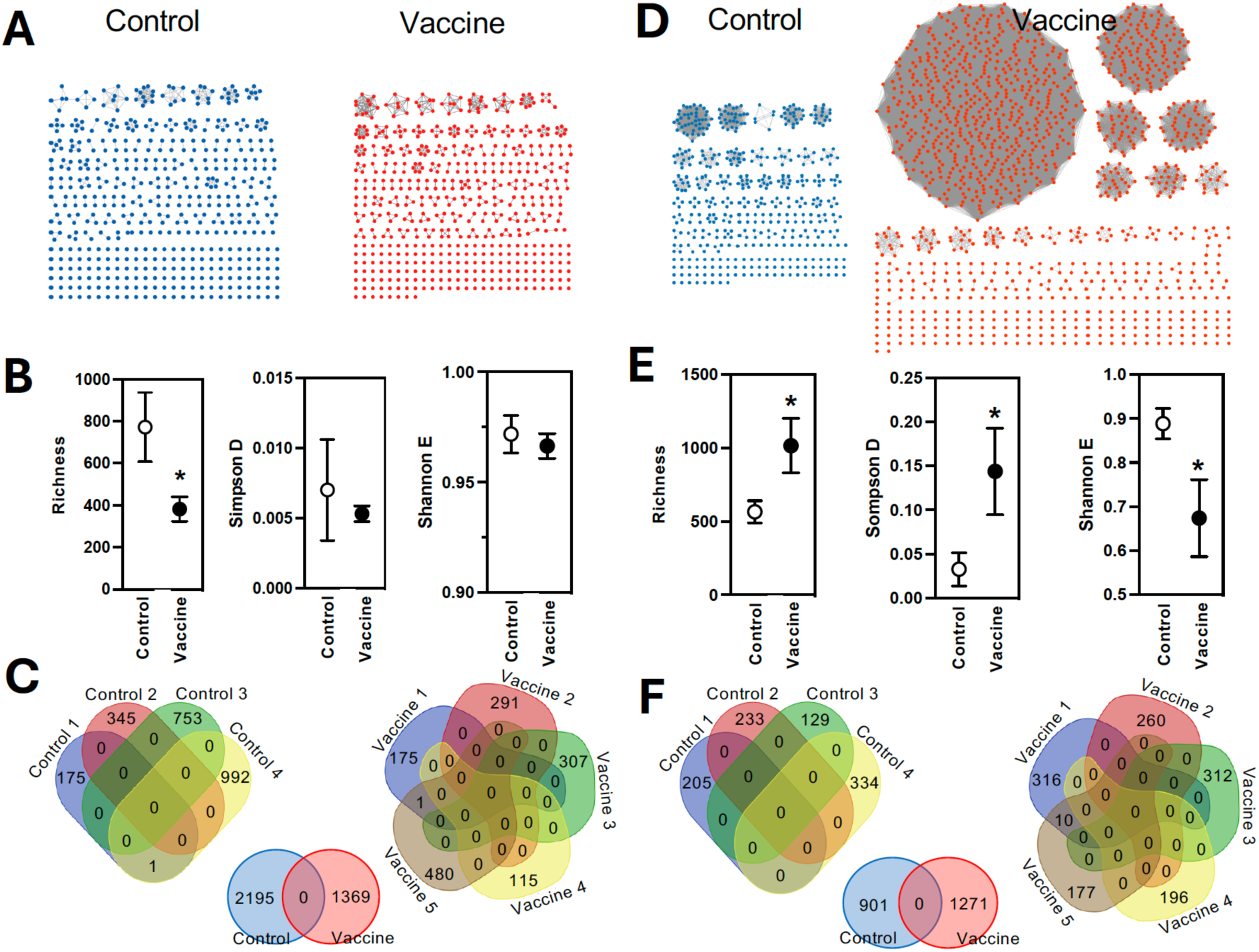
IgG and TCR repertoires of *T. cruzi* infected dogs. Representative IGH CDR3 (**A**) and TCRB CDR3 (**D**) repertoires from individual dogs. Nodes represent individual CDR3 sequences, and edges link identical sequences highlighting clonal expansion of B or T cells, respectively. IGH CDR3 (**B)** and TCRB CDR3 (**E**) Richness, Simpson and Shannon diversity indices. Data are represented as mean ± SEM of 4-5 dogs per group. *Indicates a statistically significant difference between vaccine-treated and control dogs. Ven diagrams of IgGH CDR3 (**C**) and TCRB CDR3 (**F**) sequences overlap among *T. cruzi* infected control dogs (left), among vaccine treated dogs (right), and between vaccine treated and control dogs (middle).

On the other hand, analysis of the TCRB repertoire indicated major changes in vaccinated dogs compared with untreated controls, with a strong expansion of a small number of TCRB CDR3 clonotypes together with a large increase in repertoire diversity, that resulted in broader but also more focused TCRB repertoires as (Figure 5D and E). As for IgGH CDR3 repertoire, TCRB CDR3 repertoire was highly private and no public TCRB CDR3 clonotypes were detected among PBMCs of control dogs, while in vaccinated dogs, 10/1271 (0.8%) TCRB CDR3 clonotypes were public, although shared between only two dogs (Figure 5F), and none of the TCRB CDR3 clonotypes that expanded following vaccination were present in control dogs, suggesting that these were likely induced by vaccination.

As CDR3 diversity derives from rearrangements of the respective TCR and IgG variable (V), joining (J) and diversity (D) gene segments during T and B cell maturation, we examined gene usage frequency among dogs. Analysis of IgGH V, D, J gene usage indicated that vaccination resulted in very limited changes in gene usage frequency with IGHV3-38 and IGHV3-41 together with IGHD4, IGHD3 and IGHJ4 being the most frequently used in both control and vaccinated dogs (Figure 6A). Thus, amino acid composition of the resulting IgGH CDR3 clonotypes was highly similar between groups (Figure 6B), again suggesting limited effect of vaccination on the antibody response. On the contrary, analysis of TCRB V, D, J gene usage revealed significant differences in gene usage, with TRBV18 with TRBJ2-6 and TRBJ1-4 frequent use in control dogs replaced by the use of TRBV10 and TRBV26 with TRBJ1-2 and TRBJ2-5 as some of the major changes (Figure 6C). Accordingly, amino acid composition of TCRB CDR3 clonotypes reflected these changes in gene usage and the expansion of several clonotypes (Figure 6D). Together, these results highlight important changes in TCR repertoire, with the strong expansion of novel clonotypes not present in naturally infected dogs resulting in a broader and more focused repertoire that persisted at least two months after the last vaccine dose, while the IgGH repertoire was much less affected by vaccination at this time.

**Figure 6.**
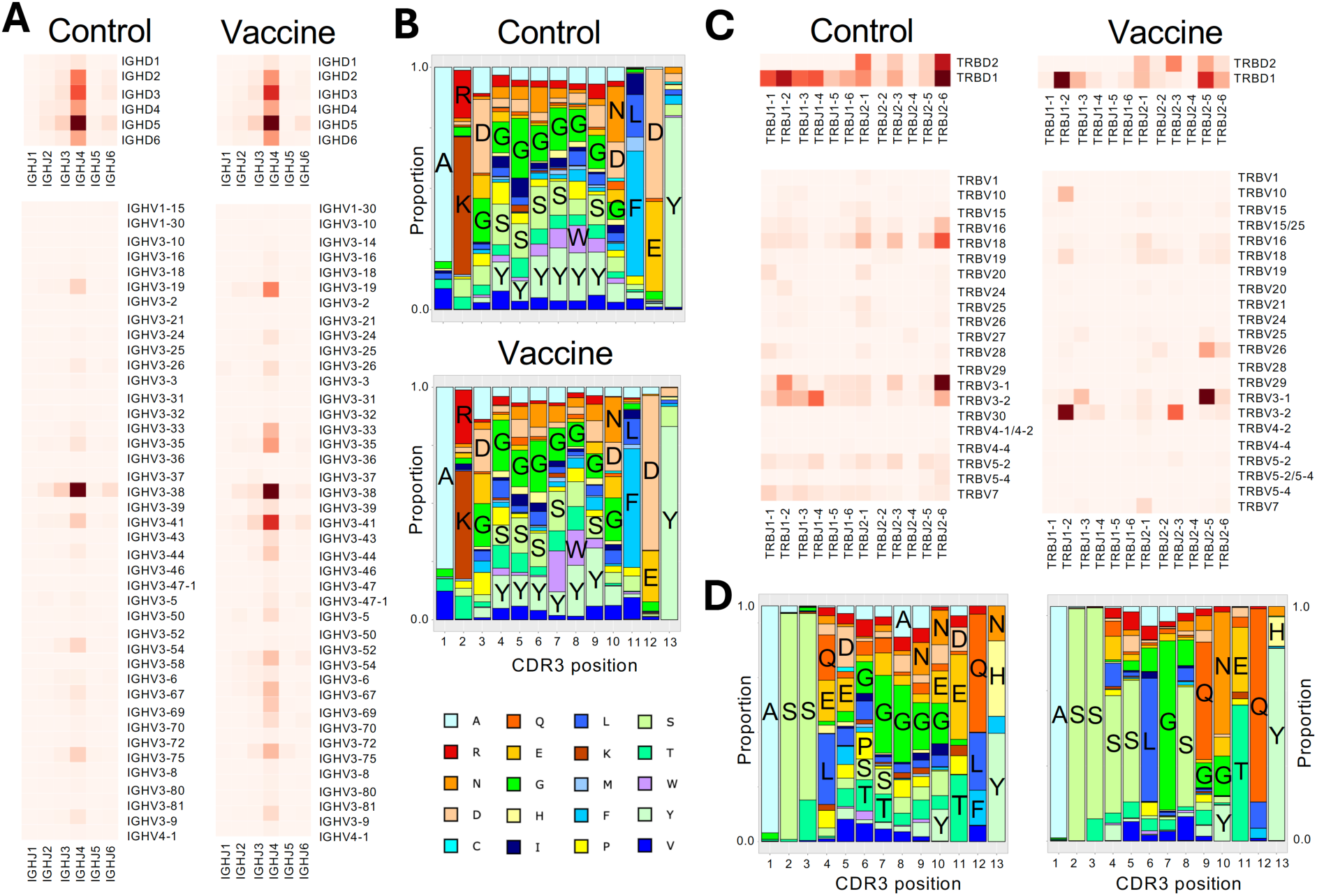
IGH and TCRB V, D, J gene usage among *T. cruzi* infected dogs. **(A)** Heatmaps of Canlupfam IGHV, IGHJ and IGHD gene usage in *T. cruzi* infected dogs (left panels) and in vaccine-treated infected dogs (right panels). (**B**) Amino acid composition of IGH CDR3 sequences from *T. cruzi* infected dogs (top panel) and in vaccine-treated infected dogs (bottom panel). (**C**) Heatmaps of Canlupfam TRBV, TRBJ and TRBD gene usage in *T. cruzi* infected dogs (left panels) and in vaccine-treated infected dogs (right panels). (**D**) Amino acid composition of TCRB CDR3 sequences from *T. cruzi* infected dogs (left panel) and in vaccine-treated infected dogs (right panel).

## DISCUSSION

A veterinary vaccine against *T. cruzi* would be a key new tool for improved Chagas disease control, allowing to reduce parasite transmission to humans ^32,54^ and improving care for companion animals and working dogs by reducing the impact of clinical disease ^10,55^. We performed here the first randomized field trial of a therapeutic vaccine against *T. cruzi* in client-owned dogs with natural infections from a rural endemic area in southern Mexico. About 27% of dogs are infected in this area, with an incidence of 6-9% per year ^37^. Infected dogs were treated with three doses of a recombinant vaccine formulated with MPLA as adjuvant, and followed for up to six months.

A first key observation was that the vaccine was safe and well tolerated, with no notable side effects during the six months follow-up, and this was confirmed by a wellness check at 12 months after vaccination. This confirms and expands previous studies in mouse models, dogs and non-human primates with experimental *T. cruzi* infection and various formulations of this vaccine ^17,28^. In terms of therapeutic efficacy, vaccination led to a reduced blood parasite burden and stopped/delayed the progression of cardiac alterations for at least six months, as assessed by qPCR and ECG profiles. Indeed, multivariate analysis of ECG profiles indicated that the ECG profile of untreated control was increasingly distinct from that of uninfected dogs, suggesting cardiac disease progression during the six months of follow-up. On the other hand, the ECG profile from vaccine-treated dogs shifted closer to that of uninfected dogs, suggesting that cardiac alterations were at least stopped/delayed during this period, although follow up for a longer time would strengthen this finding. Studies in experimentally infected mice have previously shown that a therapeutic vaccine can indeed reverse, at least in part, chronic cardiomyopathy caused by *T. cruzi* ^56^. Our data provide strong evidence of the clinical benefit of therapeutic vaccination, observed for the first time in the context of chronic natural infections with a broad diversity of parasite strains that included TcI, TcII, TcIV, TcV and TcVI ^37^. Thus, these results warrant further development of this veterinary vaccine.

Remarkably, vaccine effect on *T. cruzi* parasite burden appeared greater in dogs presenting infections with mixed DTUs, compared to those infected with TcI only. This seems in agreement with previous work suggesting that mixed infections may be less favorable for the parasite, possibly due to strain competition and/or cross-immunity limiting parasite multiplication in the host ^38^, and it suggests that mixed infections may be easier to control with therapeutic vaccination. A better understanding of how infecting strain composition, which we proposed to refer to as the “cruziome”, influences disease progression and play a role in response to drug treatments and vaccines is thus critical ^36^. Nonetheless, all vaccine treated dogs benefited from vaccine treatment as no outlier or non-responsive dog was detected, irrespective of parasite DTUs present. This is also very encouraging as the vaccine antigens used are thought to be highly conserved rather than strain-specific ^34,35^. Furthermore, although DLA typing was limited, no DLA-restriction of vaccine efficacy emerged, supporting the immunogenicity of vaccine antigens for a broad DLA genetic background. These observations also support a general use of the vaccine across dog populations in *T. cruzi* endemic areas.

Analysis of the immune response and PBMC transcriptomic profile also provided new critical insight on the mechanisms of action and correlates of vaccine efficacy. Indeed, multiple previous studies in experimentally infected mice had shown that therapeutic vaccination induced a re-orientation of the ongoing immune response, with the activation of a broad but balanced T cell response leading parasite control ^17,28^. Our results agree with this model as we identified T cell responses and the IFNγ pathway rather than antibodies as primarily affected by therapeutic vaccination and these responses were detected two months after the last vaccine dose suggesting long-term effects. In particular, we detected a strong expansion of a limited number of TCR clonotypes that resulted in a more diverse TCR repertoire after vaccination, but with several over-represented clonotypes. Accordingly, these changes in TCR repertoire were associated with important changes in TCRB V, D, J gene usage and CDR3 amino acid composition. Furthermore, both antibody and TCR repertoires were highly privates, with each dog presenting unique repertoires suggesting responses to different antigens/epitopes. Nonetheless, a few public TCR clonotypes were identified following vaccination, indicating some convergent T cell responses likely against vaccine antigens/epitopes. Indeed, as expanded TCR clonotypes were not detected prior to vaccination in *T. cruzi* infected dogs, they may have been induced by the therapeutic vaccination, although their specificity remains unclear as it was not assessed in the present study. Highly private immune repertoires of *T. cruzi* infected dogs observed here are very similar to those of naturally-infected non-human primates, which were also highly variable among individuals, irrespective of the cruziome or disease progression status ^57^. In that sense, a divide-and-conquer strategy spreading the immune response over a large number of target epitopes may reduce its effectiveness to control *T. cruzi*, while refocusing on a more limited number of epitopes that become over-represented following vaccination may help overcome this division.

In addition, our observations provide new evidence of the key role of innate immunity in *T. cruzi* pathogenesis and in vaccine efficacy, which is striking as dogs were in the chronic phase of infection when vaccinated. Indeed, transcriptomic analysis pointed to several innate processes driving inflammation in infected dogs, and vaccine treatment strongly downregulated this pro-inflammatory environment. Deconvolution of cell types suggested a particular inhibition of NK cells and monocytes by vaccination, but pathway analysis indicated the modulation of even broader innate processes also involving neutrophils, the complement system, antimicrobial peptides and reactive oxygen species. Metabolic shifts also appear to be occurring among PBMCs. These observations are consistent with therapeutic vaccination leading to a broad reprogramming of the ongoing innate response still present in the chronic phase of infection ^58–60^. Indeed, data from non-human primates previously suggested that chronic *T. cruzi* infection is associated with long term alterations of innate responses suggesting innate memory such as in trained immunity, with a pro-inflammatory environment affected by parasite strain composition ^61^. Although innate processes are often neglected and not investigated during the chronic phase of *T. cruzi* infections, a similar study in human patients also highlighted pathways associated with innate immunity correlated with the severity of chronic cardiac disease ^62^. Thus, a likely hypothesis is that innate responses in the early steps of *T. cruzi* infection lead to long-term functional programming of innate immune cells that persists during the chronic phase and may contribute to the inflammatory damage, as proposed before ^61^. Remarkably, in addition to its effects on T cell responses, therapeutic vaccination appeared to also reset/reprogram this ongoing innate memory to reduce inflammation, suggesting broader mechanisms of action and a complex interplay between innate cells and T cells ^63^. This is consistent with the generation of memory-like NK cells in response to vaccination of mice with a highly attenuated *T. cruzi* strain and their key role in protecting against a secondary infectious challenge ^64^. Lastly, several lncRNAs were found to be differentially expressed after vaccination, providing potential biomarkers of vaccine response and/or disease progression. While their function is still unknown, they may be involved in some of the immune/metabolic regulatory processes identified and such roles may be investigated in future studies.

In conclusion, this first randomized field trial of a therapeutic vaccine in dogs with natural *T. cruzi* infection showed that the vaccine was safe and effective at controlling parasites from a broad diversity of strains including TcI, TcII, TcIV, TcV and TcVI. Indeed, vaccination resulted in a significant decrease in blood parasite burden and prevented/stopped the progression of cardiac alterations assessed by ECGs, although vaccine efficacy may be higher for infections with mixtures of strains compared to simple infections. Analysis of host responses indicated that vaccine efficacy was associated with a broader TCR repertoire with a strong expansion of a few clonotypes, evidencing T cell activation sustained for at least two months following the last vaccine dose, while antibody responses were minimally affected. At the same time, therapeutic vaccination acts through a major reset/reprogramming of the ongoing innate memory to reduce inflammation. The observed clinical benefit from therapeutic vaccination in chronically infected hosts warrants the further development of a veterinary vaccine and provide support for a human therapeutic vaccine. Also, further studies should help uncover details on the interplay between innate and adaptive immunity in Chagas disease pathogenesis and vaccine efficacy.

## Acknowledgements

We thank all dog owners for their interest and participation in the study. This work was supported by grants R21AI175523 and R01AI162907 from the National Institute of Allergy and Infectious Diseases to ED. The content is solely the responsibility of the authors and does not necessarily represent the official views of the National Institutes of Health. Support was also provided through grant 557608 from the Carlos Slim Foundation via Baylor College of Medicine.

