## Supplementary tables for "Randomized field trial of a therapeutic vaccine against *Trypanosoma cruzi* natural infection in dogs and correlates for efficacy"

**Supplementary Table 1. DLA typing of dogs**

| **DLA** | **Allele** | **Vaccine** | **Control** |
| --- | --- | --- | --- |
| DLA-79 | 79*001:01 | 4 (0.4) | 3 (0.375) |
|  | 79*001:05 | 4 (0.4) | 4 (0.5) |
|  | 79*002:02 | 2 (0.2) | 1 (0.125) |
| DLA-88 | 88*002:01 | 2 (0.2) | - |
|  | 88*003:02 | 1 (0.1) | - |
|  | 88*005:01 | - | 2 (0.25) |
|  | 88*006:01 | 1 (0.1) | 3 (0.375) |
|  | 88*012:01 | 4 (0.4) | - |
|  | 88*016:03 | - | 2 (0.5) |
|  | 88*017:01 | 1 (0.1) | - |
|  | 88*501:01 | 1 (0.1) | - |
|  | 88*039:01 | - | 1 (0.125) |
| DLA-DRB1 | DRB1*001:01 |  | 2 (0.25) |
|  | DRB1*006:01 | 1 (0.1) | - |
|  | DRB1*008:02 | - | 1 (0.125) |
|  | DRB1*009:01 | 1 (0.1) | 1 (0.125) |
|  | DRB1*011:01 | 1 (0.1) | - |
|  | DRB1*013:01 | 1 (0.1) | - |
|  | DRB1*015:01 | 5 (0.5) | 2 (0.25) |
|  | DRB1:016:01 | - | 1 (0.125) |
|  | DRB1*018:01 | 1 (0.1) | - |
|  | DRB1*020:02 | - | 1 (0.125) |
| DLA-DQB1 | DQB1*002:01 | 1 (0.1) | 1 (0.125) |
|  | DQB1*003:01 | 2 (0.2) | - |
|  | DQB1*004:01 |  | 1 (0.125) |
|  | DQB1*008:01 | 2 (0.2) | 2 (0.25) |
|  | DQB1*013:02 | 2 (0.2) | - |
|  | DQB1*020:02 | 1 (0.1) | 1 (0.125) |
|  | DQB1*023:01 | - | 1 (0.125) |
|  | DQB1*026:01 | 1 (0.1) | 1 (0.125) |
|  | DQB1*034:01 | 1 (0.1) | 1 (0.125) |
| DLA-DQA1 | DQA1*001:01 | 3 (0.3) | 3 (0.375) |
|  | DQA1*002:01 | 1 (0.1) | - |
|  | DQA1*003:01 | - | 1 (0.125) |
|  | DQA1*005:01 | 1 (0.1) | - |
|  | DQA1*006:01 | 4 (0.4) | 2 (0. 25) |
|  | DQA1*009:01 | - | 1 (0.125) |
|  | DQA1*021:01 | 1 (0.1) | - |
|  | DQA1*022:01 | - | 1 (0.125) |

DLA was typed in 5 vaccinated dogs and 4 controls, corresponding to 10 and 8 alleles per group, respectively. Data are presented as N (frequency) for each allele. There were no significant differences in allele frequencies between groups, except for DLA-88 (X^2^=0.26, P=0.87 for DLA-79; X^2^=20.2, P=0.009 for DLA-88; X^2^=13.5, P=0.13 for DLA-DRB1; X^2^=8.1, P=0.42 for DLA-DQB1; X^2^=8.8, P=0.27 for DLA-DQA1 alleles).

**Supplementary Table 2. Summary of follow-up of dogs**

|  | Baseline | 3 Months | 6 Months | 12 Months |
| --- | --- | --- | --- | --- |
| Control | N=15 (100%) | N=11 (73%)  Lost to follow-up:  2 pregnancies  1 accident  1 potential distemper | N=7 (47%)  Lost to follow-up:  1 accident  1 withdrawal  1 potential distemper  1 unknown death | N=5 (33%)  Lost to follow-up:  2 unknown status |
| Vaccine | N=16 (100%) | N=16 (100%) | N=15 (94%)  Lost to follow-up:  1 pregnancy | N=11 (69%)  Lost to follow-up:  1 accident  1 unknown death  2 potential distemper |
