## Supplementary Figure for "Randomized field trial of a therapeutic vaccine against *Trypanosoma cruzi* natural infection in dogs and correlates for efficacy"

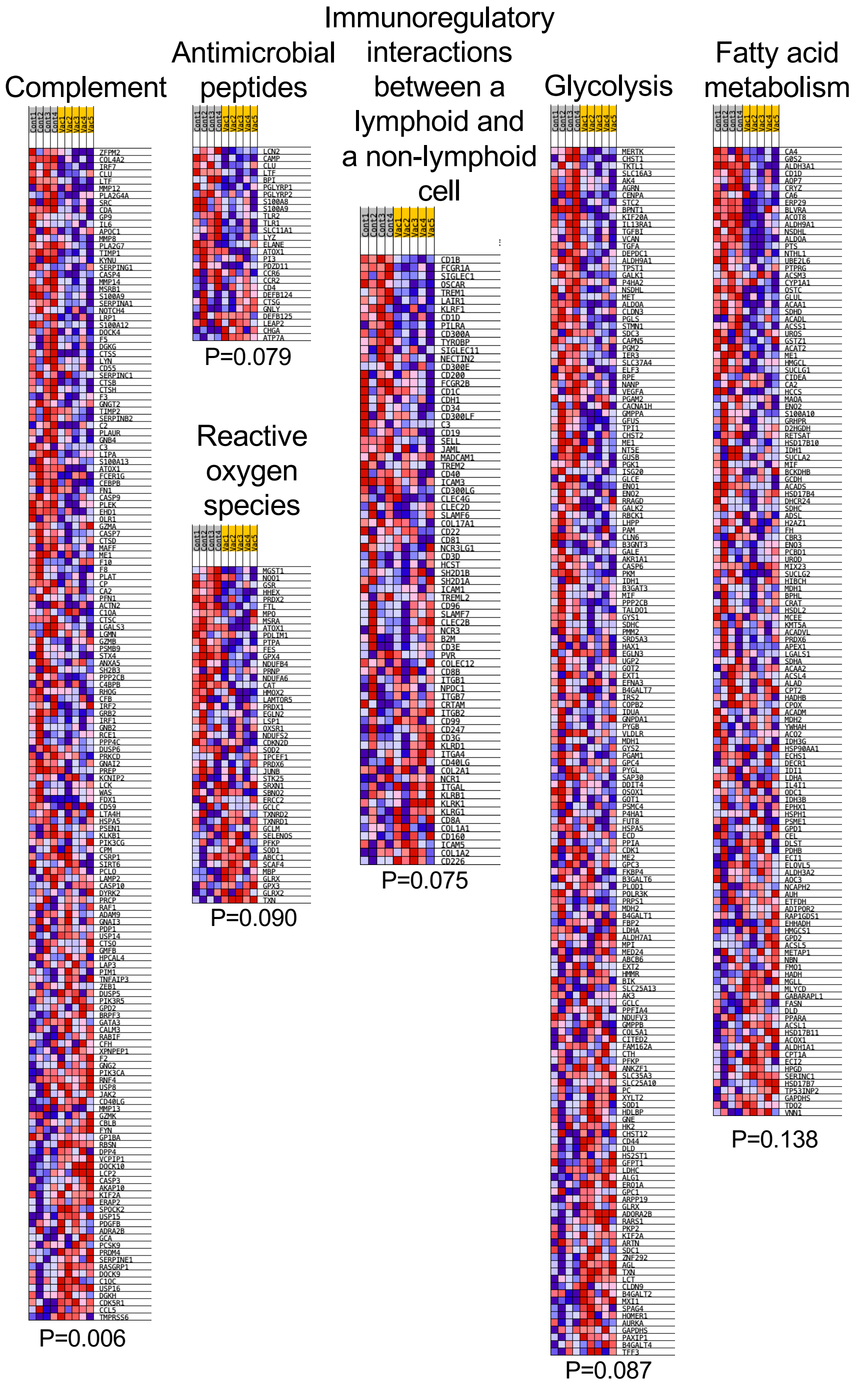


**Supplementary figure 1. GSEA pathways in vaccine-treated and control dogs.** Heatmaps of differential gene expression profiles from selected GSEA pathways among vaccine-treated (Vac) and control dogs (Cont) with *T. cruzi* infection. The statistical significance of the pathways is indicated at the bottom of each heatmap (FDR-adjusted P value).
